# The cytokine genes of *Oncorhynchus masou formosanus* include a defective interleukin-4A gene

**DOI:** 10.1101/2023.11.26.568658

**Authors:** Ying-Hsuan Yen, De Yu Zheng, Shu Yuan Yang, Jin-Chywan Gwo, Sebastian D. Fugmann

**Affiliations:** Department of Biomedical Sciences, Chang Gung University, Taoyuan, Taiwan; Graduate Institute of Biomedical Sciences, Chang Gung University, Taoyuan, Taiwan; Department of Obstetrics and Gynecology, Chang Gung Memorial Hospital, Linkou, Taiwan; Department of Aquaculture, National Taiwan Ocean University, Keelung, Taiwan; Department of Nephrology, Chang Gung Memorial Hospital, Linkou, Taiwan; Center of Molecular and Clinical Immunology, Chang Gung University, Taiwan

**Keywords:** salmonid fish, cytokines, type 2 immune response

## Abstract

*Oncorhynchus masou formosanus* (Formosa land-locked salmon) is a critically endangered salmonid fish endemic to Taiwan. To begin to understand how its drastic change in lifestyle from anadromous to exclusively river-dwelling is reflected in its immune genes, we characterized the genes encoding six cytokines (IL-2A, IL-2B, IL-4A, IL-4B1, IL-4B2, and IL-17A/F2a) important for T cell responses as no genomic data is available for this fish. Interestingly, all genes appeared homozygous indicative of a genetic bottleneck. The *IL2* and *IL17A/F2a* genes and their products are highly similar to their characterized homologs in *Oncorhynchus mykiss* (rainbow trout) and other salmonid fish. Two notable differences were observed in *IL4* family important for type 2 immune responses. First, *O. m. formosanus* carries not only one but two genes encoding IL-4B1 proteins and expansions of these genes are present in other salmonid fish. Second, the *OmfoIL4A* gene carries a 228 bp deletion that results in a premature stop codon and hence a non-functional IL-4A cytokine. This suggests a reduced ability for T cell responses against parasitic infections in this species.

## 1. Introduction

*Oncorhynchus masou formosanus* (Formosa land-locked salmon) is a critically endangered salmonid fish that is endemic to Taiwan and is threatened by farming and global warming (Gwo et al., 2019). This species arose likely after the last glacial period when the southernmost *O. masou* population became trapped in the upper reaches of the Tachia river in the central mountains of Taiwan where it is now exclusively found in the Chichiawan tributary (Healey et al., 2011). The warming water temperatures in its lower reaches had created a temperature barrier preventing migration and hence the fish acquired a non-anadromous life style. In contrast, many of the Northern *O. masou* populations whose descendants are now found in Japan are still anadromous (Morita, 2011).

Immune genes are the most rapid evolving genes (Vinkler et al., 2023) and there is an enormous level of diversity across fish not only in terms of gene sequences but also in terms of copy numbers. The evolutionary history makes *O. m. formosanus* an interesting example to study how immune genes adjust to an altered life style and environment. There is currently no sequence information available for the nuclear genome of this species or the closely related *O. masou* from Japan, and hence we set out to determine the sequences of several cytokine genes that are critical for T cell mediated immunity and known to show considerable sequence variation across vertebrates.

### 1.1 Interleukin-2 (IL-2)

After the activation of naïve T cells, their proliferation is primarily driven by interleukin-2 (IL-2) and this cytokine is therefore central to all T cell responses. In addition, IL-2 plays important roles in T_H_ and T_reg_ cell differentiation and it also promotes cell killing by natural killer cells (Spolski et al., 2018). In contrast to there being a single IL-2 in mammals, many teleost fish encode two closely related IL-2 cytokines in their genomes that both share very limited similarity at the amino acid sequence level (around 15% identity) to their single mammalian orthologues (Secombes et al., 2011). In salmonid fish, the two IL-2 encoding genes *IL2A* and *IL2B* reside in genomic regions displaying high levels of synteny (Wang et al., 2018) and are thus thought to have arisen from the salmonid-specific whole genome duplication (WGD) event (Macqueen and Johnston, 2014). Despite IL-2A and IL-2B of *Oncorhynchus mykiss* (rainbow trout) being only 42% identical to each other at the amino acid level, there are no gross differences in terms of *IL2A* and *IL2B* expression and their ability to stimulate T cells (Wang et al., 2018). While they appear to signal through different receptors (Wang et al., 2018), it remains unclear at this point whether they are functionally redundant.

### 1.2 Interleukin-4 (IL-4)

IL-4, together with the related IL-13, is the defining cytokine during type 2 immune responses of T_H_2 cells against extracellular parasites in mammals (Lloyd and Snelgrove, 2018). The first fish IL-4 discovered in *Tetraodon nigrovirdis* has only 11-16% identity to mammalian IL-4 and IL-13 (Li et al., 2007). However, in spite of the very low sequence conservation, this cytokine and its gene loci are conserved throughout vertebrates. *Lepisosteus oculatus* (spotted gar), a basal non-teleost ray-finned fish belonging to a lineage that diverged before the teleost WGD, still carries the ancestral locus with a single copy of a *IL4*/*IL13* gene (hereafter referred to as *IL4* for simplicity) that is flanked by *KIF3A* and *RAD50* (Wang and Secombes, 2015a). A tandem gene duplication in the lineage from which mammals arose gave rise to a *IL4* and *IL13* gene pair that is still flanked by *KIF3A* and *RAD50* (Wang et al., 2016). In constrast, the teleost WGD and the subsequent salmonid WGD gave rise to four distinct but syntenic *IL4* gene loci in *O. mykiss* and many other salmonid fish: *IL4A* and *IL4p* (a pseudo gene) both neighboring *RAD50* (or remnants thereof, respectively), and *IL4B1* and *IL4B2* both neighboring *KIF3As* (Wang et al., 2016). A comprehensive study of the expression patterns of *O. mykiss* IL4s indicated that *IL4A* has a higher basal expression in all tissues, whereas *IL4B1* and *IL4B2* transcript levels are low under basal conditions but highly inducible during immune responses (Wang et al., 2016). Surprisingly, the abilities of recombinant IL-4A and IL-4Bs to stimulate immune cells only differed in the kinetics but not by the magnitude of the responses, and thus the distinct biological roles of the individual cytokines still remain to be elucidated.

### 1.3 Interleukin IL-17

IL-17s are among the most ancient cytokines that can be traced back to *C. elegans* (Chen et al., 2017), and the mammalian IL-17 cytokine family has six members: IL-17A, IL-17B, IL-17C, IL-17D, IL-17E, and IL-17F. Its most prominent member IL-17A (frequently simply referred to as IL-17) is the signature cytokine of T helper 17 (T_H_17) cells (Mills, 2023). This cytokine, together with IL-17F, triggers anti-microbial responses against extracellular pathogens by inducing the production of antimicrobial peptides by immune cells, fibroblasts, epithelial cells and other non-immune cells (McGeachy et al., 2019). It also initiates the production of neutrophil-attracting chemokines. The other IL-17s are also produced by innate immune cells and non-immune cells in the context of innate immune responses (Mills, 2023). The IL-17 repertoire of teleost fish, and even more so of salmonid fish, is expanded compared to that of mammals due to one (or two) additional WGDs. Notably, however, IL-17E appears to be absent from all teleost genomes. Teleost fish express three IL-17s with homology to IL-17A and IL-17F that commonly are referred to as IL-17A/F1, IL-17A/F2, and IL-17A/F3 (Okamura et al., 2023). *O. mykiss* and *Salmo salar* (Atlantic salmon) carry duplicated *IL17A/F1* genes (*F1a* and *F1b*), duplicated *IL17A/F2* genes (*F2a* and *F2b*), and single copies of genes encoding IL-17A/F3 and a more distantly related factor named IL-17N (Wang et al., 2015) that was first discovered in *Takifugu rubripes* (Korenaga et al., 2010). The steady-state expression levels of these genes vary across tissues with no clear pattern emerging (Wang et al., 2015). Similarly, the broad stimulation of immune cells from with synthetic pathogen-associated molecular patterns or recombinant cytokines resulted in surprisingly low changes in the expression levels of any of the *IL17A/Fs* and *IL17N* (Wang et al., 2015). Hence the unique roles of each of the paralogues in immune responses are still unclear.

To begin to assess the evolution of these important cytokine families in *O. m. formosanus*, we set out to amplify, clone and sequence the gene loci for *IL2A*, *IL2B*, *IL4A*, *IL4B1*, *IL4B2*, and *IL17A/F2a* and we now report the sequence of all six genes. Two notable differences were observed compared to other salmonid fish: a deletion that renders the classical *IL4A* gene non-functional and an apparent duplication of the *IL4B1* gene.

## 2. Material and Methods

### 2.1 Oligonucleotides

All oligonucleotides (Suppl. Table S1) were synthesized by a commercial provider (Genomics, Taiwan).

### 2.2 Genomic DNA

As *O. m. formosanus* is an endangered species, we decided to minimize disturbance to this species in the wild and executed this study with genomic DNA that was isolated from pre-existing ethanol preserved tissue samples (obtained by Prof. JC Gwo). An ethanol preserved fin tissue sample from a single individual #848 (male, body length 305 mm, weight 290 g) was minced with a scalpel, and the EasyPure Genomic DNA Spin Kit (Bioman) was then used to purify the gDNA according to the manufacturer’s protocol. *S. salar* genomic DNA was purified from a tissue sample obtained from a local sushi restaurant.

### 2.3 PCR

For each gene two separate PCR reactions were set up using Phusion polymerase (ThermoScientific), the primer pairs listed in Suppl. Table S1, and genomic DNA as the template. The PCR program to amplify the *OmfoIL17A/F2a* genomic fragment (IL17 –OmIL17AgF5, OmIL17AgR1) started with an initial denaturation step at 98°C for 2 min followed by 35 cycles of 98°C for 20 sec, annealing at 54°C for 20 sec, and extension at 72°C for 30 sec; a final extension step 72°C for 7min concluded the reaction. To amplify the genomic locus of *O. m. formosanus IL2A*, *IL2B*, *IL4A*, *IL4B1*, and *IL4B2* the extension time was increased to 1 min.

### 2.4 Cloning and sequencing of PCR products

All PCR products were gel-purified using the EasyPure PCR/Gel Extraction Kit (Bioman) according to the manufacturers protocol, ligated into pJET1.2 vector using the CloneJET PCR Cloning Kit (ThermoScientific), and transformed into DH5α *E. coli*. Plasmids were purified using the Tools Plasmid Mini Kit (Biotools), positive clones were identified by restriction digests and their sequences were determined by a commercial Sanger sequencing service (Genomics, Taiwan).

### 2.5 Bioinformatics

The sequences of the *O. m. formosanus* cytokine genes were aligned with the corresponding *O. mykiss* gene loci sequences using BLAST (NCBI). Exon/intron boundaries were identified and annotated manually. Multiple sequence alignments and phylogenetic tree calculations were conducted using CLUSTALW (Larkin et al., 2007) and MEGA 11 (Tamura et al., 2021).

### 2.6 Sequence information

All annotated gene sequences and the respective transcript and protein sequences were obtained from NCBI Genbank (<u>ncbi.nlm.nih.gov</u>) and Ensembl (www.ensembl.org). Currently unannotated genes from species for which a complete genome sequence was available were manually annotated using annotated homologs as the template. All sequence data and the information about their source is provided in an excel sheet as Supplementary Data. The Genbank accession numbers for the *O. m. formosanus* genes are OR228408 (*OmfoIL2A*), OR228409 (*OmfoIL2B*), OR228410 (*OmfoIL17A/F2a*), OR228411 (*OmfoIL4A*), OR228412 (*OmfoIL4B1a*), OR228413 (*OmfoIL4B1b*), OR228414 (*OmfoIL4B2*).

## 3. Results

### 3.1 Cloning of cytokine gene loci

For each of the six cytokine genes, we aligned the genomic sequences of four salmonid fish (*O. mykiss*, *O. nerka*, *O. keta*, and *S. salar*) and designed primers pairs (Suppl. Table S1) in regions that showed 100% sequence conservation such that the forward primer hybridizes upstream of the respective ATG translation start and the reverse primer downstream of the stop codons, respectively. To clone the *O. m. formosanus* homologs of these genes, these primer pairs were then used in PCR reactions with the genomic DNA of one adult individual as the template. For five of the cytokines we obtained PCR products that closely matched in size those expected other salmonid fish (Fig. 1), but for *OmfoIL4A* we only obtained a smaller product of about 1.0 kb compared to the predicted 1.2 kb based on all other species (described and discussed in detail below).

**Figure 1.**
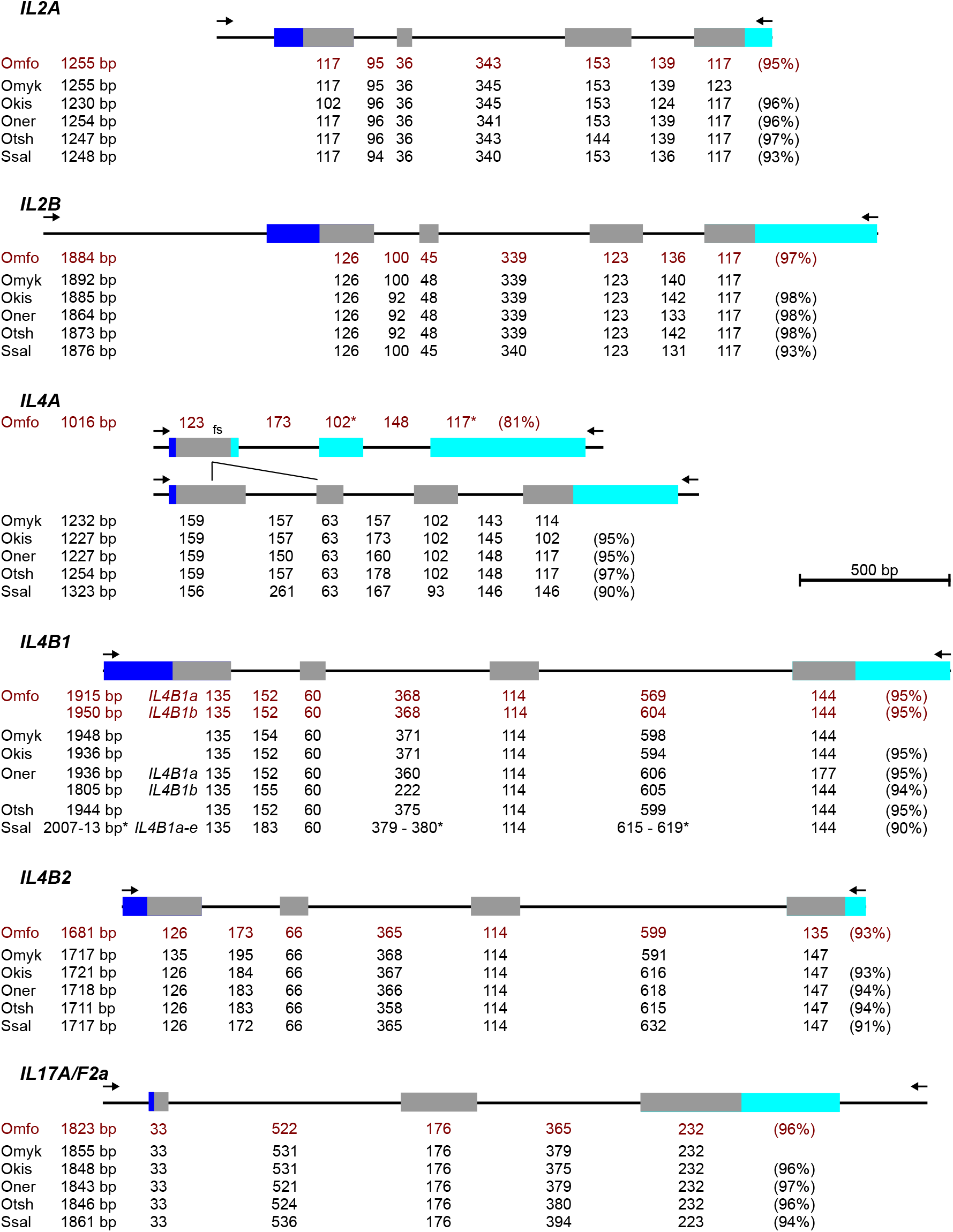
Schematic representation of the cytokine gene loci of *O. m. formosanus* and related salmonid fish. The coding region (grey), 5’UTR (dark blue), and 3’UTR (light blue) are indicated as boxes. The total length of the PCR products (arrows represent the primers) obtained from *O. m. formosanus* (Omfo) genomic DNA and the predicted sizes of the products from other species (*Oncorhynchus mykiss* [Omyk], *Oncorhynchus kisutch* [Okis], *Oncorhynchus nerka* [Oner], *Oncorhynchus tshawytscha* [Otsh], and *Salmo salar* [Ssal]) are shown as well as the lengths of the exons (only the coding sequence) and introns. The percentage of identity at the nucleotide level compared to the *O. mykiss* genes are listed on the right. For the *OmfoIL4A* gene the position of the deletion and the resulting frame-shift mutation (fs) relative to the *OmykIL4A* gene are indicated. Note that *S. salar* harbors five predicted *IL4B1* genes (*IL4B1a-e*) in its genome and hence a range of sizes (*) is listed, whereas for *O. nerka* and *O.m. formosanus* the sizes for *IL4B1a* and *IL4B1b* are provided individually.

All PCR products were subsequently cloned and sequenced. Multiple sequence alignments revealed two highly similar (96.5%) but distinct sets of nucleotide sequences for the cloned *IL4B1* PCR products (hereafter referred to as *OmfoIL4B1a* and *OmfoIL4B1b*, respectively), while for all other cytokine genes the sequences of all clones were identical. This apparent lack of genetic variation suggests that the genetic diversity across the small extant population is low. We then annotated the *O. m. formosanus IL2A*, *IL2B*, *IL4A*, *IL4B1a, IL4B1b*, *IL4B2*, *IL17A/F2a* gene loci guided by the annotations in other salmonid fish (Fig. 1). For the 5’ end of the first exons (transcription start) and the 3’ ends of the last exons (polyadenylation sites) the respective *O. mykiss* genes were used as reference points except for *IL2B* for which the *O. nerka* gene served as the template. It is important to note, that the transcription starts and poly-adenylation sites for each cytokine are not fully consistent across the public genome datasets as many current annotations rely on computational gene prediction models rather than comprehensive transcriptome sequence data.

For all *O. m. formosanus* cytokine genes but *IL4A* the overall gene structure is highly conserved with respect to the number of exons and introns, their lengths, and their sequences (Fig. 1). Furthermore, all donor/acceptor splice sites conform to the GT/AG consensus. For *IL4A*, however, comparison with the *OmykIL4A* locus revealed a 228 bp deletion encompassing the 3’-end of exon 1 and intron 1 resulting in an *OmfoIL4A* gene with only three exons (Fig. 1); in contrast, the IL4A genes of all other salmonid fish closely resemble that of *O. mykiss*.

To confirm that this deletion is in fact present in the genome and not a cloning or PCR artefact caused by repeats or secondary structures, we conducted PCRs with a primer pair (Omfo-IL4AF1/R1) immediately flanking the deletion (Fig. 2A). We compared the product obtained for the *O. m. formosanus* sample to that of a *S. salar* genomic DNA, a salmonid fish with an intact *IL4A* gene. The former gave rise to a 323 bp amplicon whereas the *S. salar* product was much larger matching the expected size of 550-650 bp for the intact *IL4A* gene loci of salmonid fish (Fig. 2B). This confirms that the deletion is indeed present in at least one allele of the *OmfoIL4A* locus. Due to the shorter size this allele could be preferentially amplified by PCR. To assess whether there is possibly also a second intact allele, an additional control PCR was conducted combining the original reverse primer (O-IL4AR) and a new forward primer (O-IL4AFi1) hybridizing at the beginning of intron 1 that is 100% conserved across at least four salmonid fish (Fig. 2A). Here a robust PCR product of the expected size of about 1100 bp was only observed for *S. salar* while no such product could be amplified from *O. m. formosanus* genomic DNA (Fig. 2C). This indicates that the deletion is indeed present on both alleles and hence an actual feature of the *OmfoIL4A* locus.

**Figure 2.**
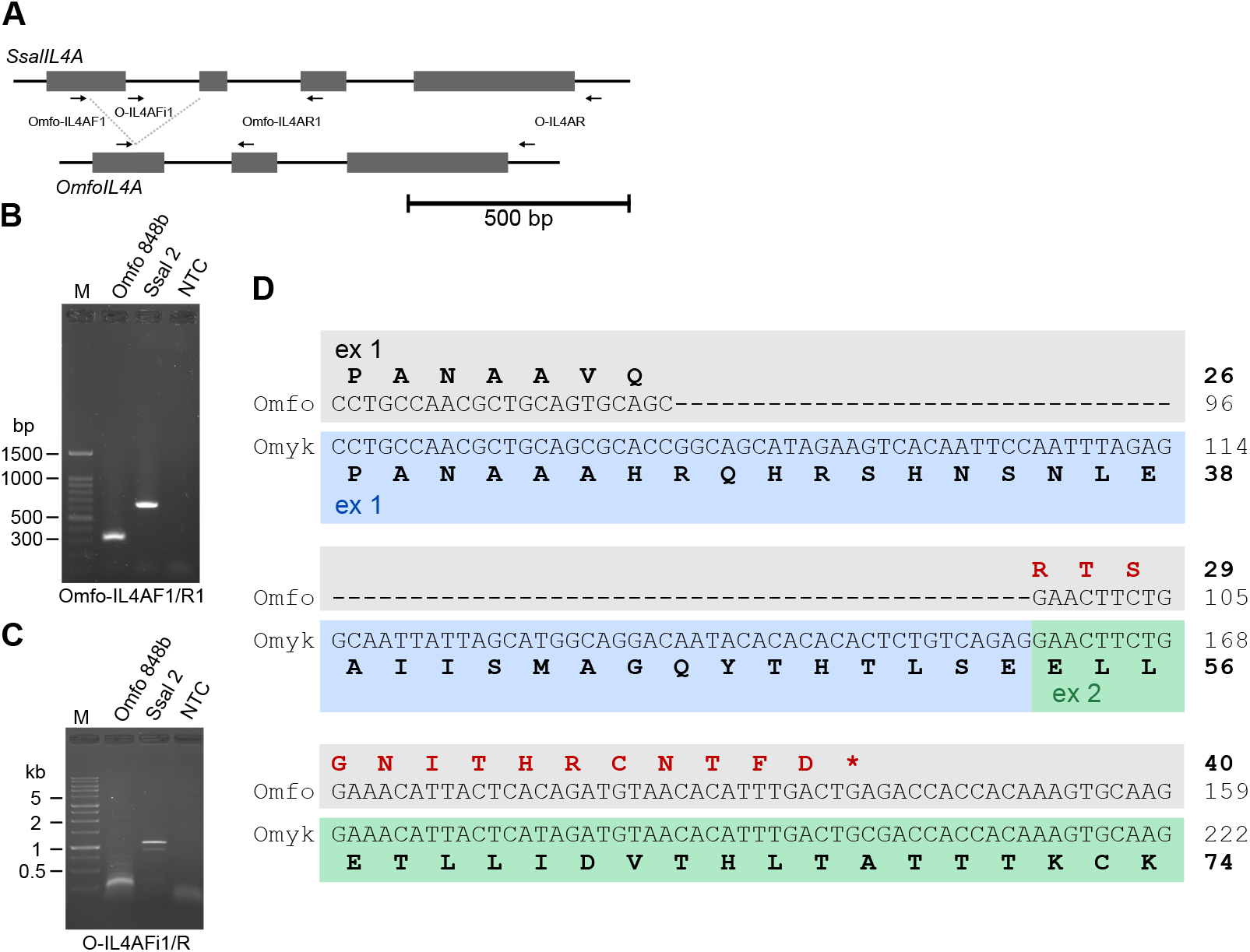
A deletion in the *IL4A* gene of *O. m. formosanus*. Schematic comparison of the *IL4A* gene loci in the *S. salar* and *O. m. formosanus* genomes. Exons are shown as boxes, primer locations are indicated by arrows, and the deletion within the *OmfoIL4A* gene is illustrated by dotted lines. (**B**+**C**) PCR reaction using the indicated primer pairs were conducted using genomic DNA from one individual of each species (Omfo 848b and Ssal 2) as templates with a non-template control (NTC) serving as a negative control. Products were resolved on agarose gels with a marker for size comparison. In (**B**) products of the expected sizes (Omfo: 323 bp, Ssal: 655 bp) were observed for both, whereas in (**C**) only the Ssal template gave rise to a respective band of the expected size of 1122 bp (note the small band for Omfo 848b corresponds to a primer dimer).

### 3.2 Comparison of cytokine protein sequences

Based on our gene annotation we derived the predicted mRNA and corresponding amino acid sequences of all the encoded cytokines. The resulting proteins, with the exception of the predicted *OmfoIL4A* gene product (discussed in detail below), share strong similarity with those from other salmonid fish (Fig. 3) with no overt differences that could likely alter their functions. Moreover, the cysteines involved in the disulfide bonds required for structural integrity are conserved (data not shown). This includes four cysteines typical for the IL-4 cytokines (Wang and Secombes, 2015b), four cysteines and two serines in the cysteine knot-like structure of IL-17s (Hymowitz et al., 2001), and the two and three pairs of cysteines in the IL-2A/IL-2B of salmonid fish (Wang et al., 2018).

**Figure 3.**
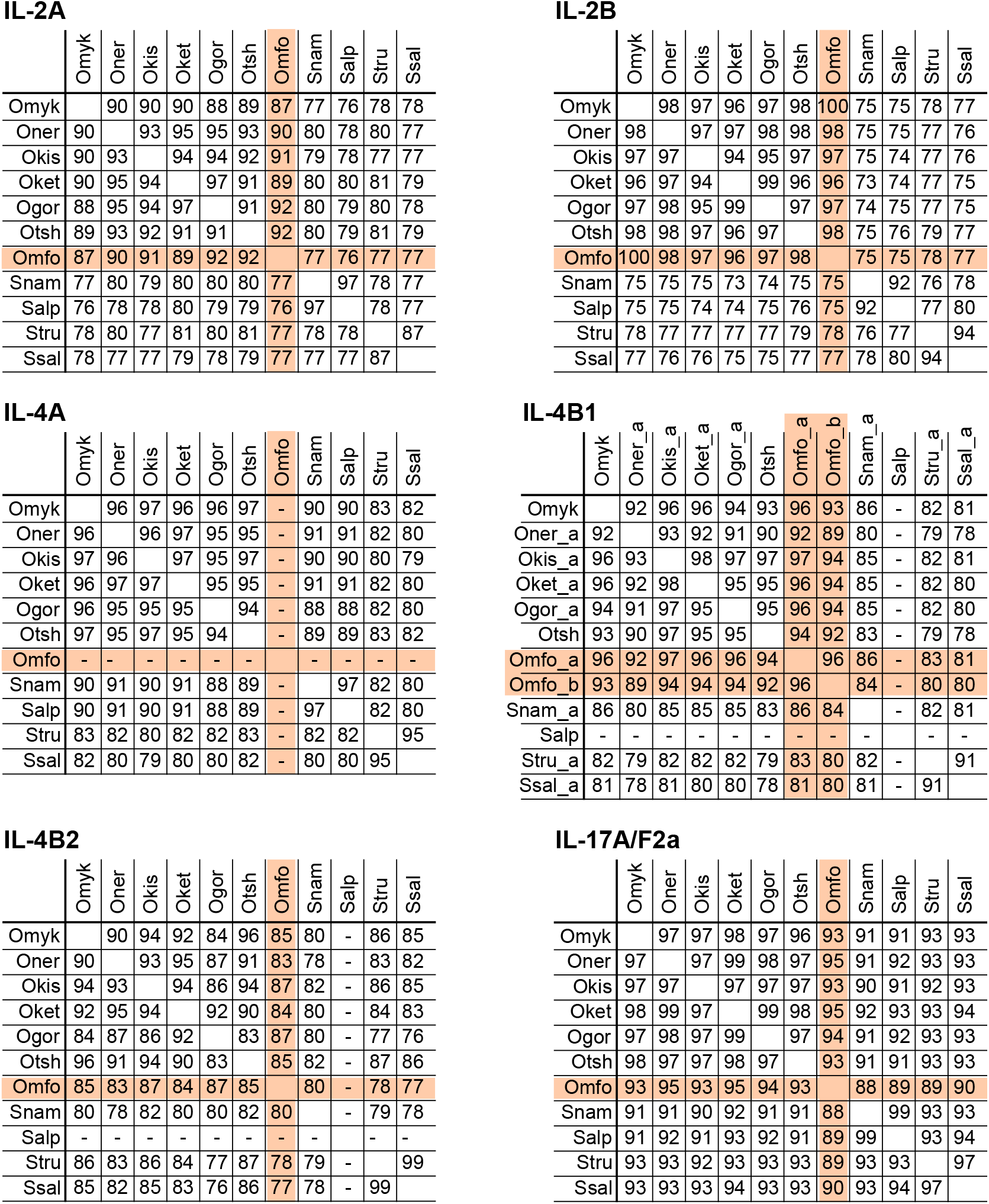
Sequence similarity of salmonid fish cytokines. The amino acid sequences of the indicated cytokines from salmonid fish were aligned and the pairwise identities were calculated using CLUSTALW. The values for *O. m. formosanus* are highlighted in red. The *OmfoIL4A*, *SalpIL4B1*, and *SalpIL4B2* genes carry premature stop codons and hence were excluded (-) from the alignments. Note that for IL-4B1 only the products of one representative gene of all species (except for *O. m. formosanus*) were used.

To determine their evolutionary relationships, we then calculated phylogenetic trees including the respective cytokines from ten additional salmonid fish and the *Esox lucius* (Northern pike) proteins serving as the outgroup (Figs. 4, 5, and 6). This fish diverged from the Salmonidae before the WGD and carries only single copies of the *IL2*, *IL4A*, *IL4B*, and *IL17A/F2* (Rondeau et al., 2014) gene loci. As expected, the IL-2As and IL-2Bs split into separate clusters (Fig. 4), as do the IL-4A, IL-4B1 and IL-4B2 sequences (Fig. 5). In addition, within each of these clusters, the Atlantic salmon (*S. salar* and *Salmo trutta*), and char (*Salvelinus alpinus* and *Salvelinus namaycush*) form distinct subgroups that diverge from the base of the Pacific salmon species (*Oncorhynchus*), expect for the IL-4B2 cluster where the Atlantic salmon clusters resides in the middle of their Pacific cousins. In line with the phylogenetic relationship between *E. lucius* and salmonid fish, ElucIL-4A and ElucIL-4B reside at the base of the IL-4A and IL-4B1/2 branches. Within the *Oncorhynchus* species, the *O. m. formosanus* cytokines are in their expected position at or near the base of the clusters for IL-2A and IL-2B as well as for IL-4B1 and IL-4B2, but their relative positions within each cluster vary likely due to the overall low level of sequence divergence. In contrast, OmfoIL-17AF2a groups together with its *Oncorhynchus nerka* homolog. Overall, the phylogenetic trees reflect the established evolutionary relationships.

**Figure 4.**
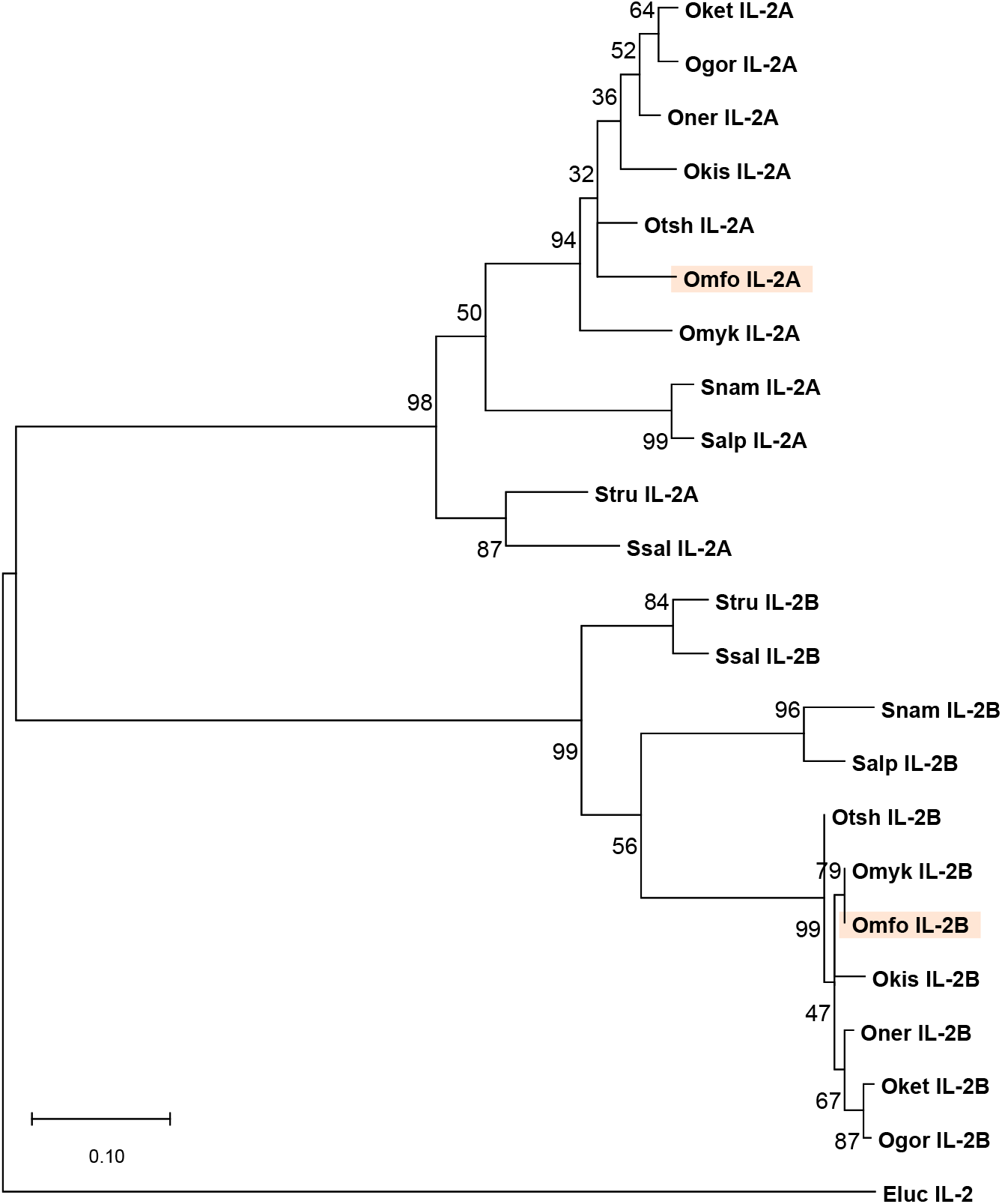
Phylogenetic tree of salmonid fish IL-2s. The evolutionary relationships of the two IL-2s from *O. m. formosanus* (IL-2A and IL-2B, reported in this manuscript) and of IL-2s from a range of other salmonid fish were inferred by using the Maximum Likelihood method and JTT matrix-based model in MEGA 11. The sole IL-2 from *Esox lucius* served as an outgroup. The tree with the highest log likelihood (−2179.54) is shown. The percentage of trees in which the associated taxa clustered together is shown next to the branches.

**Figure 5.**
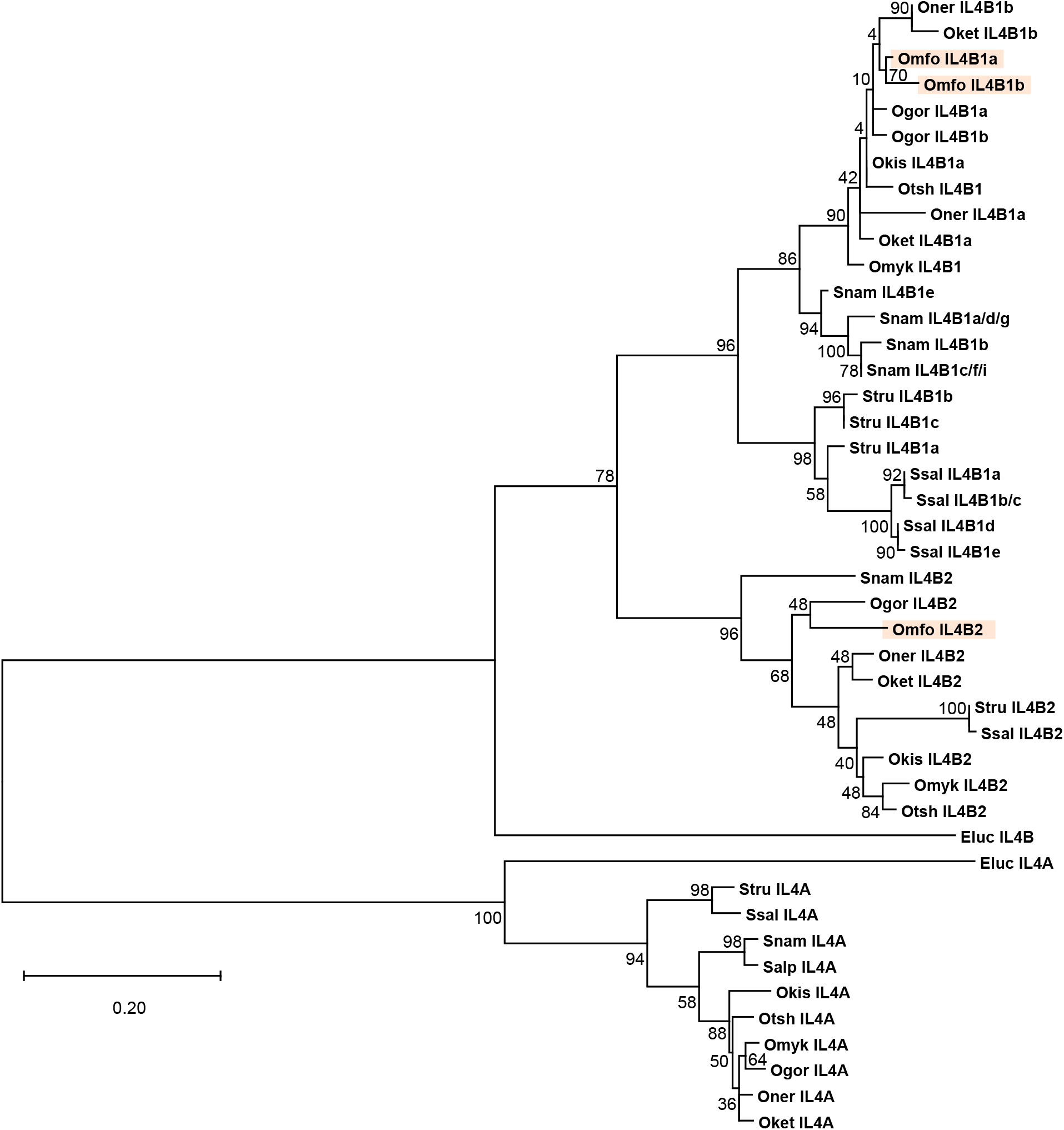
Phylogenetic tree of salmonid fish IL-4s. The evolutionary relationships of the three IL-4s from *O. m. formosanus* (IL-4B1a, IL-4B1b, IL-4B2) and the IL-4s from a range of other salmonid fish were inferred by using the Maximum Likelihood method and JTT matrix-based model in MEGA 11. IL-4A and IL-4B from *Esox lucius* served as an outgroup. Note that gene products of *OmfoIL4A*, *Salp-IL4B1*, and *SalpIL4B2* with a predicted premature stop codon were excluded from the analysis. The tree with the highest log likelihood (−3207.65) is shown. The percentage of trees in which the associated taxa clustered together is shown next to the branches.

**Figure 6.**
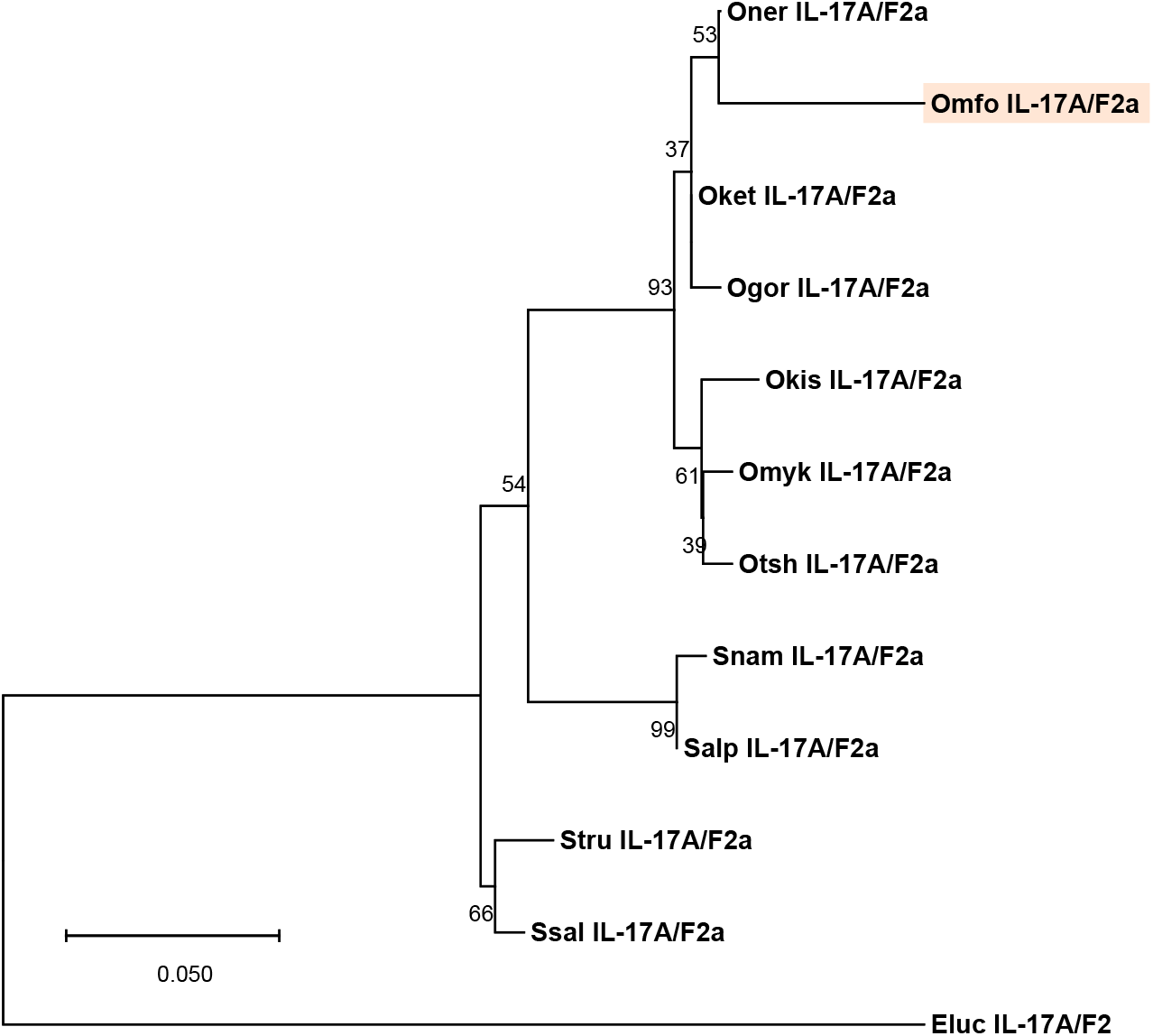
Phylogenetic tree of salmonid fish IL-17A/F2a. The evolutionary relationships of IL-17A/F2a from *O. m. formosanus* and a range of other salmonid fish were inferred by using the Maximum Likelihood method and JTT matrix-based model in MEGA 11. IL-17A/F2 from *Esox lucius* served as an outgroup. The tree with the highest log likelihood (−848.89) is shown. The percentage of trees in which the associated taxa clustered together is shown next to the branches.

Among *Oncorhynchus*, the branch lengths for the IL-2Bs are clearly shorter than those calculated for the IL-2A (Fig. 4), an observation consistent with the higher overall similarity of the IL-2Bs (94-100% identity) compared to their IL-2A paralogs (87-97% identity) (Fig. 3). A comparable trend is also apparent in the IL-4s with IL-4A (94-97% identity, excluding the non-functional OmfoIL-4A) (Fig. 5) showing a higher level of conservation compared to IL-4B1 (89-98% identity) and IL-4B2(83-96% identity) (Fig. 3).

Importantly, many salmonid fish genes harbor more than one *IL4B1* gene, with *S. namaycush* carrying a total of eight functional genes (*SnamIL4B1a-i*) encoding a total of four distinct SnamIL4B1 proteins (SnamIL-4B1a/d/g, SnamIL-4B1b, SnamIL-4B1c/f/i, and SnamIL-4B1e) that can be distinguished based on their amino acid sequences. Within the phylogenetic tree the IL-4B1s from each species cluster together apart from *O. nerka* and *O. keta*, for whom the respective IL-4B1a and IL-4B1b group together. This suggests that (1) the *IL4B1* loci are less stable than the *IL4B2* loci with gene expansions largely occurring independently in each species, but (2) a single common gene duplication preceded the split of *O. nerka* and *O. keta*.

### 3.3 The non-functional OmfoIL4A gene product

Based on the genomic sequence the predicted *OmfoIL4A* transcript is not only shorter than that of all other salmonid fish, the deletion of parts of exon 1 and intron 1 (Fig. 2A) also results in a frameshift such that it only encodes a short peptide of 40 amino acids (Fig. 2D). Moreover, only the first 27 amino acids (23 of which encode the signal peptide) share similarity to the IL-4A cytokine of *O. mykiss* and all other salmonid fish, indicating that the truncated OmfoIL4A, even if it were expressed, is unlikely to act as an IL-4 family cytokine. This strongly suggests that IL4 dependent type 2 immune responses are defective in *O. m. formosanus*.

## 4. Discussion

We set out to clone and sequence *O. m. formosanus* genes representing three important cytokine families for cellular immunity: IL-2, IL-4, and IL17. As members of all three families were found to be present in the genome, it indicates that T cell mediated immunity is overall intact. The lack of a functional *IL4A* gene, encoding one of the three IL-4 orthologues in salmonid fish, suggests that T_H_2 mediated immune responses are altered in *O. m. formosanus*. This is supported by the recent discovery of expression differences between the *IL4A* (basal) and *IL4B1/ IL4B2* (highly inducible) genes in two disease models in *O. mykiss* (Wang et al., 2016). This is likely caused by the distinct transcriptional control elements responsible for *IL4* gene regulation that got split up between the *IL4A* and *IL4B1*/*IL4B2* gene loci. In mammals *IL4* control elements reside within the neighboring *KIF3* and *Rad50* genes, and in salmonid fish these got separated between the orthologues with *KIF3s* being linked to *IL4B1* and *IL4B2* and the single *RAD50* being linked to *IL4A*. Combined with the fact that IL-4A and IL-4Bs induced not only common but also some distinct innate immune effectors and other cytokines relevant for type-2 immunity (Wang et al., 2016), this indicates that the salmonid IL4 orthologues are not entirely redundant. Hence the lack of any of them, like here in *O. m. formosanus*, will likely result in reduced type-2 immune responses. No comprehensive studies have been conducted assessing whether this species is more susceptible to extracellular pathogens or harbors a higher parasite load; obtaining regulatory approval for future *in vivo* experiments to assess such questions with this critically endangered species will be challenging. Among all salmonid fish for which genome data is available, this deletion in *IL4A* is unique to *O. m. formosanus* and thus it likely occurred after the split of the *O. masou* lineage from all other fish in the *Oncorhynchus* genus. Analysis of the respective genes of all Japanese *O. masou* subspecies will be important to inform us whether this variant is a unique genetic feature of the Taiwanese salmon.

Our sequence analysis also revealed that *O. m. formosanus* likely harbors gene loci encoding two closely related IL-4B1 proteins, IL-4B1a and IL-4B1b, sharing 96% identity at the amino acid level. Comparing these gene loci revealed 31 single nt differences, two single nt indels, and one larger 33 nt insertion that are together spread over 2 kb. As we did not discover any evidence for allelic variations in the other cytokine genes, we concluded that *IL4B1a* and *IL4B1b* likely represent two closely related but distinct genes. An analysis of the respective gene loci from other salmonid fish revealed similar patterns. While *S. alpinus* only carries a highly mutated non-functional *IL4B1* pseudogene (Suppl Fig. S2), the genomes of many salmonid fish appear to harbor more than one *IL4B1* gene (some of which are mutated pseudogenes) within the same chromosomal location (Supplementary Data file). The largest number is found in the *S. namaycush* genome with a total of eight functional *IL4B1* genes in three loci. Systematic combined RT-PCR and genomic PCR studies will be required to determine how many of those *IL4B1* genes represent genome assembly artefacts (e.g. allelic variants of a single gene appear as two separate loci) and similarly for *O. m. formosanus* only the completion of the genome will provide the final answer.

Finally, the sequence analysis of our PCR products did not reveal allelic variations like SNPs in any of the six gene loci reported here and the same pattern is seen in all other gene loci that we have analyzed thus far. As it is unlikely that all cytokine genes in *O. m. formosanus* reside on a sex chromosome, this finding indicates that the current population in the wild (from which the samples in this study were originally obtained) shows a high level of homozygosity. This supports a previous report measuring heterozygosity using microsatellite DNA loci (Yamamoto et al., 2020). The presence of a broad repertoire of haplotypes is commonly thought to be beneficial especially in genes involved in immune responses. As the extant fish population appears to rely on a limited gene pool, it raises concerns about the long-term survival of the species in the context of imminent environmental changes including the microbiota in the mountain streams of Taiwan.

## Acknowledgements

We thank Lukas A. Fugmann for help in preparing genomic DNA, and all members of the Fugmann and Yang labs for helpful discussions. This work was supported by grants of Chang Gung University [UMRPD1N0121] to S.D.F, the Chang Gung Memorial Hospital [CMRPD1M0461] and NSTC [112-2311-B-182-003-MY3] to S.Y.Y. as well as private funds from the Fugmann/Yang family.

---

**Suppl. Fig. 1.**
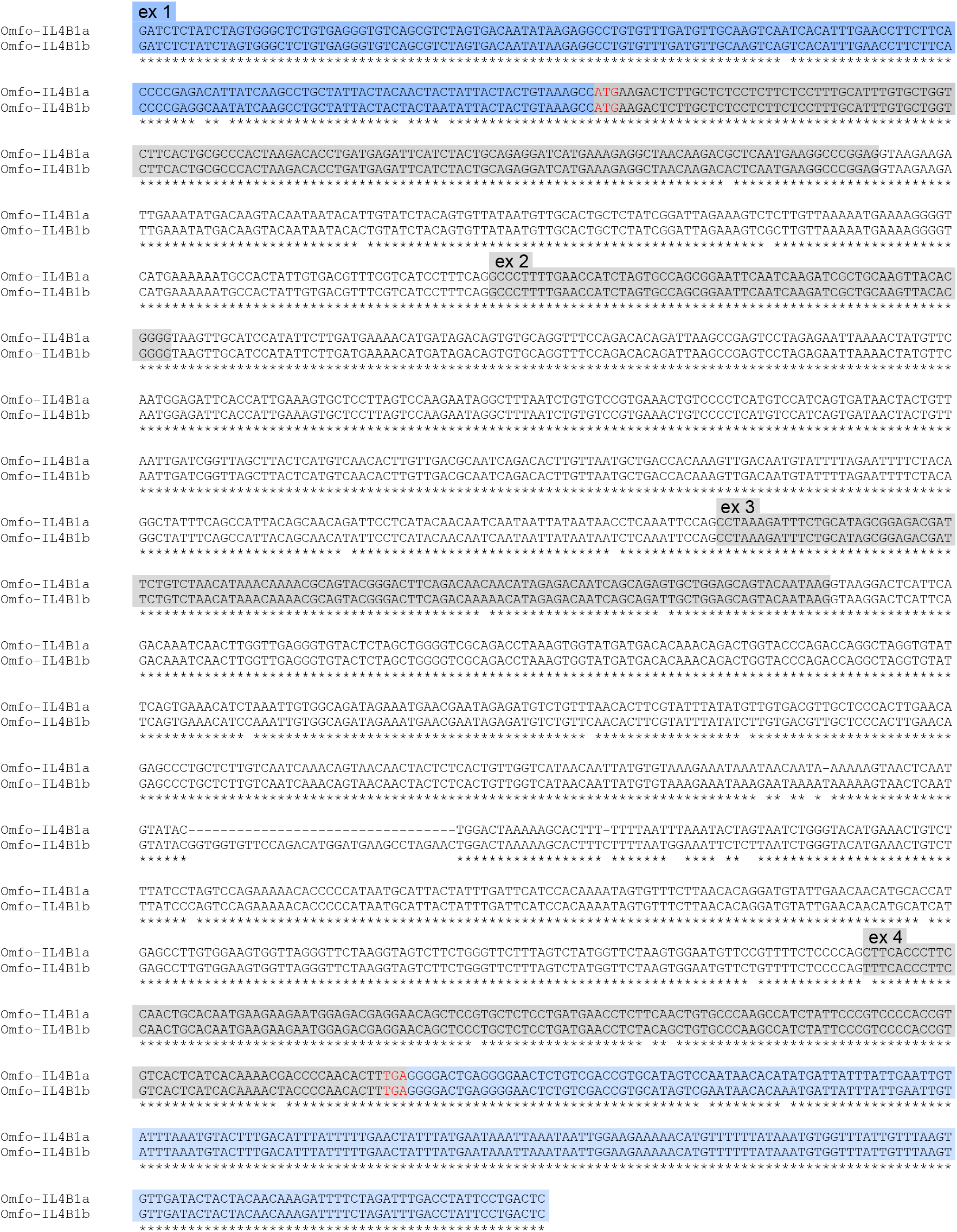
Sequence alignment of the *O. m. formosanus IL4B1a* and *IL4B1b* genes. The sequences of the *OmfoIL4B1a* and *OmfoIL4B1b* genes were aligned using CLUSTALW. Exons are indicated as boxes with UTRs in blue and coding sequences in grey, respectively. The start (ATG) and stop (TGA) codons are highlighted in red. Asterisks indicate sequence identity.

**Suppl. Fig. 2.**
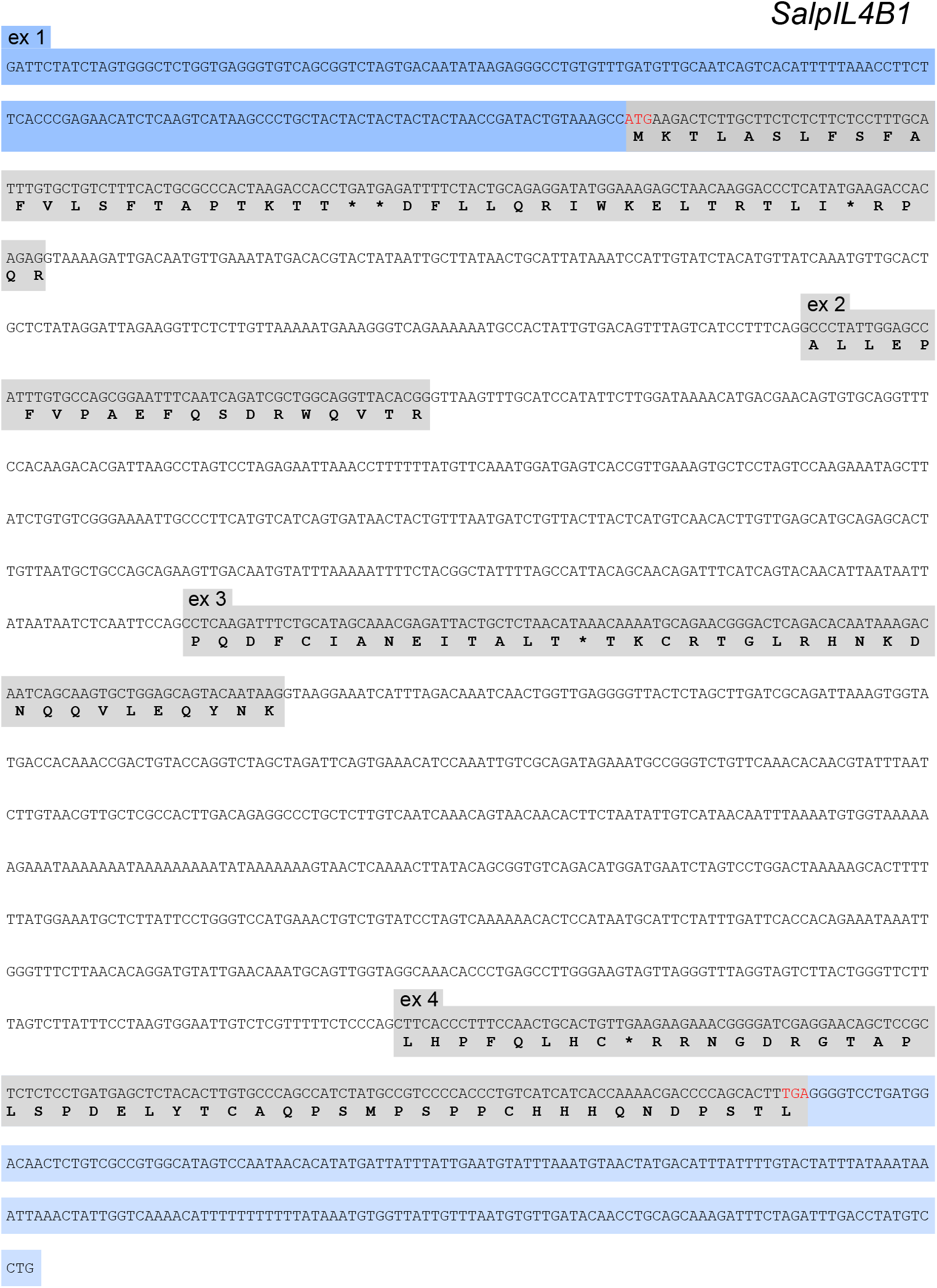
Nucleotide sequence of the *Salvelinus alpinus IL4B1* gene. Exons are indicated as boxes with UTRs in blue and coding sequences in grey, respectively. Note that the UTRs likely extend beyond the regions shown. The start (ATG) and stop (TGA) codons are highlighted in red. The translation products are shown underneath the coding sequences with asterisks indicating premature stop codons. Note that the translation products and the end of the coding sequence in exon 4 are based on the exon phases and translations present in all other salmonid fish.

**Supplementary Table 1.**
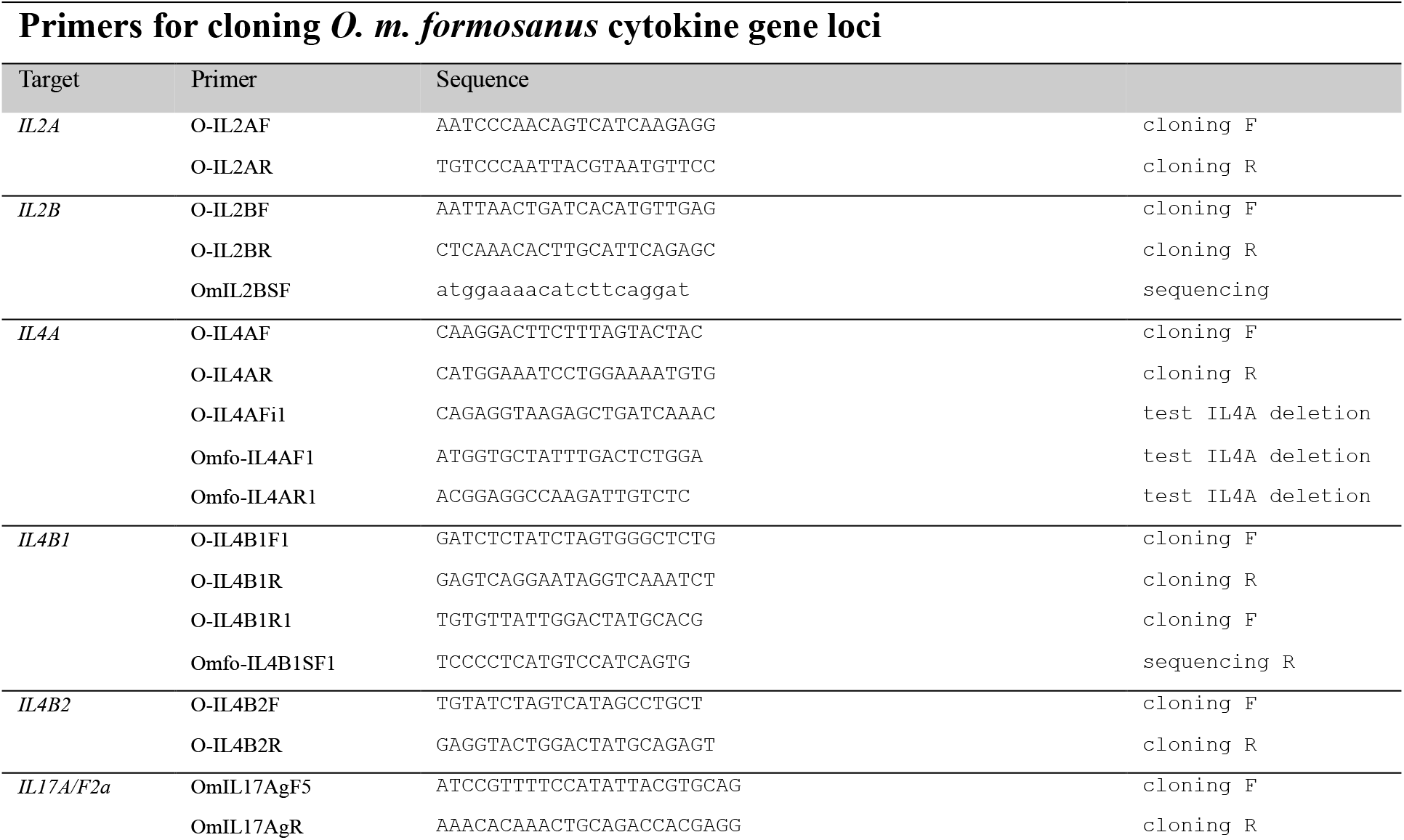

